# The Rational Irrational: Better Learners Show Stronger Frequency Heuristics

**DOI:** 10.1101/2025.09.18.676999

**Authors:** Mianzhi Hu, Darrell A. Worthy

## Abstract

Does favoring less valuable options that deliver more frequent rewards reflect flawed decision-making or an adaptive strategy under complex environments? Frequency effects, defined as a bias toward more frequently rewarded but less valuable options, have traditionally been viewed as maladaptive decision-making deficits. In the present study, we used a within-subject design in which participants completed a four-option reinforcement learning task twice, once under a baseline condition and once with a reward frequency manipulation, to test whether better baseline learning predicts greater or lesser susceptibility to frequency-based biases. Participants were first trained on two fixed option pairs and then transferred their knowledge to novel pairings in a testing phase. Across conditions, higher training accuracy generally predicted higher test accuracy, with one critical exception: on trials where a more valuable option was pitted against a more frequently rewarded but less valuable alternative, participants with higher training accuracy exhibited a stronger bias toward the more frequent option. Moreover, baseline optimal choice rates in these specific trials were unrelated to—and even slightly negatively correlated with—optimal choice rates under the frequency condition. Computational modeling further showed that participants with better baseline learning performance were better fit by frequency-sensitive models in the frequency condition and they weighed frequency-based processing more heavily than value-based processing. Overall, these findings suggest that frequency effects, rather than signaling flawed learning, manifest more strongly in individuals with better baseline learning performance. This seemingly irrational bias may, under conditions of uncertainty, reflect a flexible, adaptive strategy that emerges among the best learners when value-based approaches are costly or unreliable.

**Author Summary:** In daily life, people often face choices between familiar, frequently encountered options and unfamiliar alternatives that may be more valuable. For example, we may keep visiting a local restaurant we know well instead of trying a new one with better reviews. This tendency, known as the frequency effect, reflects a bias toward options that yield more frequent rewards, even when those rewards are smaller and suboptimal overall. Traditionally, such behavior has been interpreted as a sign of neuropsychological impairments or flawed learning, while our study found the opposite. We asked 495 participants to complete a reinforcement learning task under two conditions: one with balanced reward frequencies and another in which one option was rewarded more frequently despite being less valuable than its alternative. Surprisingly, we found that better learners in the balanced condition were more likely to show frequency effects when reward frequencies were manipulated and uneven. Computational modeling confirmed that these individuals shifted from value-based strategies to frequency-based ones when the environment made value-based decisions more difficult. These findings suggest that frequency effects are not simply errors. Instead, they may represent an adaptive shortcut that emerges more strongly in better decision-makers as a flexible strategy for navigating uncertain environments when value-based calculations are costly or unreliable

## Introduction

Would you choose to dine at a decent local restaurant you know well, or take a chance on a new one with a higher Yelp rating? Would you stick with your usual route to work, or try a newly discovered shortcut that might be faster? Would you keep using a familiar app, or switch to an unfamiliar one that promises better features? If you tend to choose the former option, you are demonstrating what cognitive psychologists call a reward-frequency-based bias, or frequency effect (1,2). Frequency effect refers to people’s tendency to favor options that have yielded more frequent rewards, even if they offer lower average long-term payoffs (1–3). This tendency can push people towards suboptimal decisions, such as dining at worse restaurants, taking longer commutes, or settling for less efficient tools. Yet under real-world constraints, where value-based calculations can be overly costly or practically impossible, sticking with familiar, reliably rewarding options may, in fact, serve as an adaptive heuristic (4). The current study seeks to examine whether such frequency effects reflect flawed value learning or instead represent a fundamental component of adaptive decision-making, thereby emerging more strongly in individuals with stronger learning and value integration skills.

Frequency monitoring underlies many classic behavioral decision-making heuristics (4)—such as tallying (5), fluency (6), and mere-exposure effects (7,8). It has been posited as one of the most basic ways the human brain encodes statistical information (9–11). In the context of reinforcement learning (RL), however, a bias toward more frequently rewarded options despite their lower overall value has been traditionally interpreted as a decision-making deficit, particularly associated with severe psychiatric conditions (12,13) or brain injuries (14,15). Supporting this view, frequency effects have been found to be more pronounced in populations with reduced cognitive functioning, including older adults (16), individuals with substance use disorders (17,18), and those with developmental disorders (19,20).

Yet more recent findings challenge this deficit-based interpretation by showing that healthy participants without neuropsychological impairments would nevertheless show frequency effects and favor the more frequently rewarded option over objectively more valuable alternatives (21–25). Moreover, these effects do not reliably distinguish between healthy and clinical populations (22,26–28). Replications across several paradigms, including the Iowa Gambling Task (IGT) (21,22,25,28), a modified version of IGT (24,29), the Soochow Gambling Task (SGT) (28,30,31), and other four-option RL tasks (1–3), suggest that reward frequency may represent a fundamental component of value learning rather than a pathological byproduct.

Why, then, do people show frequency effects? From the perspective of bounded rationality, humans make decisions under constraints of time, information, and cognitive resources, which often obscure the true structure of the environment (4,32). Under such conditions, complex value-based computations may need to give way to simpler heuristics to achieve an effort-accuracy balance. In some cases, these heuristics can even lead to better decision outcomes compared to strictly value-based strategies (32,33). Consistent with this view, research has shown that these mental shortcuts are especially favored when the decision-making environment is complex (34–36), time is limited (37,38), or information is incomplete (39,40). Frequency effects may reflect one such adaptive shortcut. Previous studies have found that frequency effects emerge primarily when value differences between options are small, whereas when differences are large and clear, reinforcement frequency exerts little influence on people’s choices (1,2). A recent study (3) further demonstrates that preference for the more frequently rewarded option increases proportionally with the level of environmental uncertainty. When outcome variance (i.e., uncertainty) was low, participants preferred the more valuable option; when variance was moderate, no clear preference emerged; and when variance was high, the frequently rewarded option was favored. Together, these findings suggest that frequency effects may function as an adaptive, supplementary strategy that becomes increasingly dominant as value-based estimations grow more difficult.

Despite their adaptive appeal though, frequency-driven decisions often lead to suboptimal performance and outcomes in laboratory tasks and potentially in real-world contexts (41,42). What remains unclear is how an individual’s baseline learning and decision-making capacity shapes susceptibility to these effects in uncertain, complex environments. One possibility is that better learners, while still influenced by reward frequency, are more resilient to frequency-based biases and therefore will rely more consistently on value-based strategies. Another possibility is that, because switching to frequency-based processing can itself be adaptive, better learners are more likely to employ this strategy and thus exhibit stronger frequency effects.

To our knowledge, no study has systematically tested these competing possibilities. The present work addresses this gap using a within-subject design in which participants completed the same RL task twice, with and without a reward frequency manipulation. We aimed to examine whether individuals who performed better during training and in the baseline condition were more or less likely to shift toward frequency-based processing when reward frequency was manipulated, leveraging RL computational modeling. Based on prior research (3), we hypothesized that switching to frequency-based processing is adaptive. Thus, we predicted that participants who demonstrated stronger learning performance would be more likely to adopt frequency-based strategies and exhibit stronger frequency effects in the frequency condition.

## Methods

### Task

The task was adapted from the four-option RL paradigm used in Hu et al. (3). Participants were presented with four options (A, B, C, D), but on each trial, they selected from a pair of only two options. The expected values (EVs) were ranked as C (0.75) > A (0.65) > B (0.35) > D (0.25). The task consisted of two phases: a training phase and a testing phase.

During the 120-trial training phase, participants repeatedly selected from only two fixed option pairs: AB and CD. After each choice, they received feedback showing the number of virtual points earned for that trial, along with a running total of their cumulative points (*Figure 1*). Reward outcomes were randomly drawn from normal distributions centered on each option’s EV, with variance approximating binomial distributions. *Table 1* shows the specific reward structure. This level of reward variance has been shown to reliably elicit frequency effects when reward frequency is manipulated (2,3). Within each training pair, one option (i.e., A in AB and C in CD) is designed to yield significantly higher rewards than the other (i.e., B and D), making it the dominant option in its pair. However, the value difference between parallel options across pairs (e.g., A vs. C or B vs. D) is much smaller, which would allow frequency effects to emerge.

**Figure 1.**
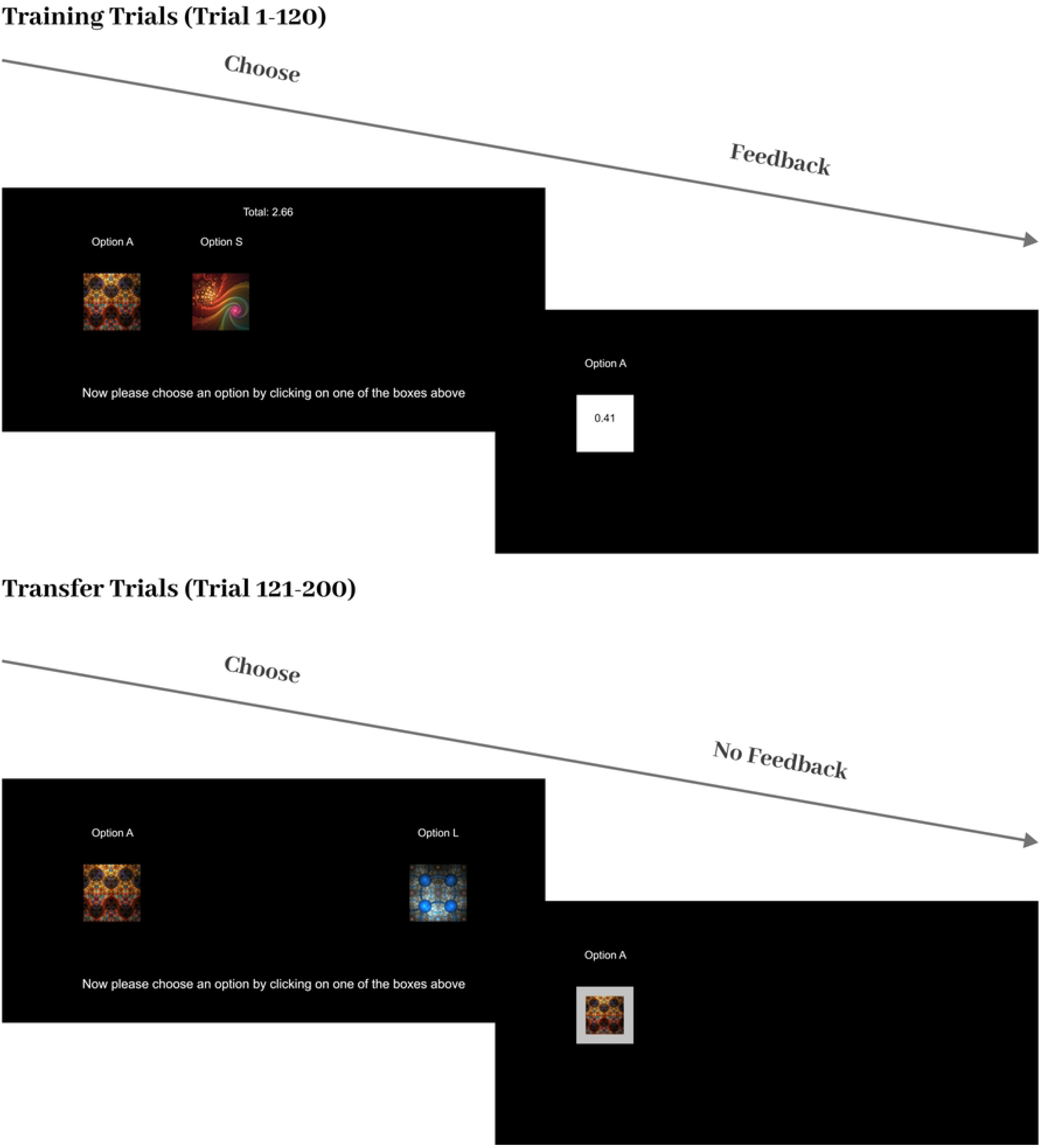
Example Trial Sequence. *Example Trial Sequences.* During the first 120 training trials, participants selected from either the two left options or the two right options (i.e., AB or CD). Option labels were randomized (e.g., Option S could correspond to A, B, C, or D in the reward schedule). The total virtual points earned were displayed at the top of the screen. After each selection, participants received feedback on the number of points gained. In the subsequent 80 testing trials, participants chose from the remaining four option combinations (i.e., 20 trials for each)—CA, BD, AD, and CB— without any cumulative point displays or feedback. They are told that the options have remained the same, but they need to make their selections based on what they learned about each option during the training phase.

**Table 1.**
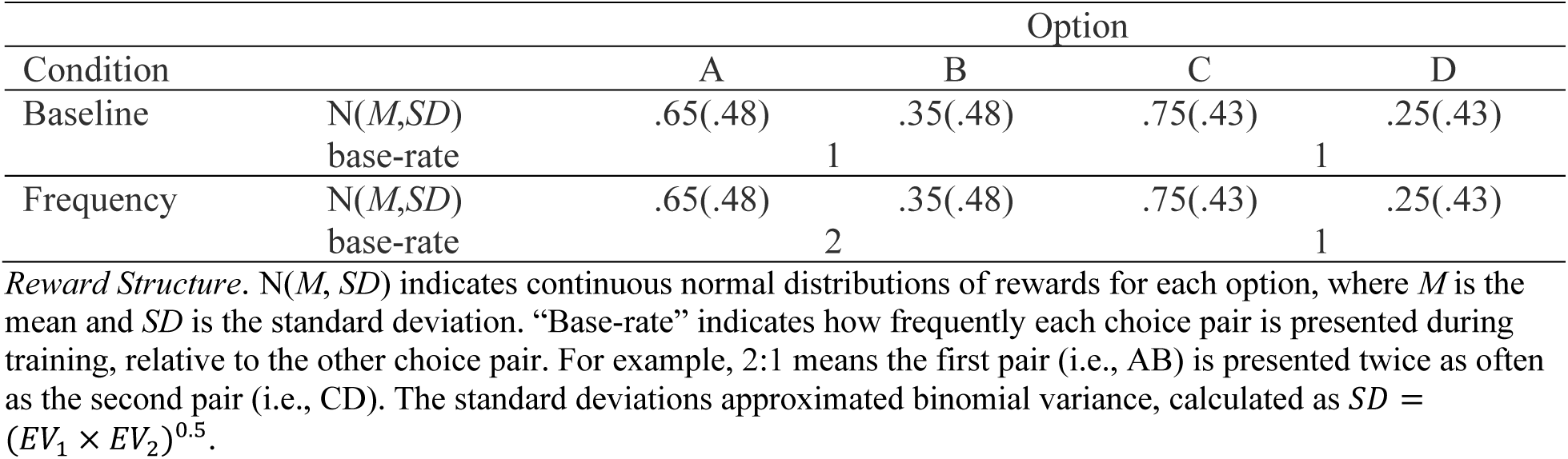
Reward Structure.

After the training phase, participants proceeded to the testing phase, where they must transfer their knowledge and select from the remaining novel pairings (i.e., AD, BD, CA, CB) without feedback. This phase had 80 trials in total, with 20 trials for each pair type.

We employed a within-subject design that manipulated the frequency of training pairs. Each participant completed the task twice under two different conditions. In the baseline condition, AB and CD pairs appeared equally often during training (60 trials each), while in the frequency condition, AB pairs appeared twice as often as CD pairs (80 vs. 40 trials), making A more frequently rewarded than C despite its lower EV. We consider CA trials as critical trials because our frequency manipulation creates a direct conflict in the CA pairing, where the slightly less rewarding option, A, is rewarded more frequently than the more rewarding option, C, in the frequency condition. Prior studies have shown that this base-rate manipulation encourages a preference shift from C to A, demonstrating frequency effects (1–3).

Participants were instructed to maximize their cumulative points by learning which options were most rewarding. Choice stimuli were four fractal images randomly drawn from a pool of 12 fractal images, with on-screen positions and image assignments randomized. While A–B and C–D always appeared together as pairs, their placement on the screen varied across participants (e.g., ABCD, CDAB, BADC).

### Participants

This study was approved by Texas A&M University Institutional Review Board (STUDY2024-1012). A priori power analysis indicated that 449 participants would provide 95% power to detect a weak-to-moderate effect size in paired samples (*d_z_* = 0.2) at an alpha level of .01 (43). Based on previous studies in our lab, we anticipated that some participants might show little evidence of learning during training, respond inattentively, or be lost due to technical errors. To account for this, we planned to recruit approximately 500 participants. In total, 501 undergraduate students from Texas A&M University participated, all of whom provided informed consent and received partial course credit for their participation. Six participants were excluded for inattentive responding, defined as consistently selecting the same option throughout the entire training session for both training pairs (e.g., always choosing A in AB and C in CD). A final sample of 495 participants went into data analysis.

The mean age was 18.93 years (*SD* = 1.01). The sample consisted of 345 females, 148 males, 1 participant who self-identified as “other”, and 1 who preferred not to answer. Racial distribution was as follows: 370 White, 60 Asian, 31 multiracial, 12 Black or African American, 3 American Indian or Alaska Native, 1 Native Hawaiian or Other Pacific Islander, and 9 who preferred not to answer. With respect to ethnicity, 135 participants identified as Hispanic or Latino, 9 preferred not to answer, and the remainder identified as non-Hispanic.

### Procedures

The experiment was administered online. Participants voluntarily enrolled through the Texas A&M University Psychology Department’s research participant pool and received partial course credit for their participation. Upon enrollment, participants were directed to the study via a secure link hosted on the Texas A&M University JATOS server, where the experiment would launch automatically.

The task was developed using the jsPsych JavaScript library for creating online behavioral experiments. Participants completed the RL task twice, once under each condition (i.e., baseline and frequency). After completing the first task, they were directed to fill out a demographic questionnaire and a series of additional surveys included as part of a separate pilot project. They were then returned to complete the second condition of the main task.

To minimize carryover effects, participants were explicitly instructed that the reward structures were completely different between the two sessions, despite the structural similarity of the tasks, and that prior knowledge should not be transferred between sessions. The order of conditions was randomized and counterbalanced across participants: 252 participants completed the frequency condition first, and 243 completed the baseline condition first. Order effects did not significantly influence behavioral patterns (see *Supplementary Table 1* and *Supplementary Figure 1*). After completing both sessions, participants were debriefed, thanked, and informed that they would receive their course credit shortly. The study concluded upon clicking the final submission button.

### Data analysis

Behavioral data were analyzed using mixed-effects models implemented via the *lme4* package in *R*. Random intercepts were included at the participant level to account for individual differences given our within-subject design. For analyses involving the percentage of optimal choices, we used the *lmer* function. For trial-level logistic regression models predicting the probability of selecting the optimal option on each trial, we used the *glmer* function.

### Computational Models

Beyond traditional behavioral analyses, we employed computational modeling to further disentangle participants’ use and weighting of distinct decision-making strategies. We focused on two major classes of reinforcement learning rules: Delta and Decay rules. For each class, we included a basic model and two widely used extensions, yielding six individual models (2×3). To capture the relative contribution of different strategies, we also constructed a seventh hybrid model by combining the best-fitting variant from each class, resulting in a total of 7 formal models.

#### Delta Model

The basic Delta rule model (44) is one of the most widely used value-based RL models. It updates the EV by incorporating the prediction error between the EV from the last trial and the actual reward received in the current trial. *EV_t+1_* for option *j* is defined as:

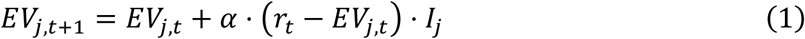

where *I_j_* is an indicator term set to 1 if option *j* is chosen on trial *t*, and 0 otherwise; *r_t_* is the reward value; and *α* is the recency learning parameter, *⍺⍺* ∈ (0,1), with higher *α* indicating greater weighting of most recent outcomes. When *α* = 0, EVs remain unchanged regardless of new outcomes; when *α* = 1, *EV_j_* is equivalent to the reward received for option *j* on its most recent selection. In this model, no memory of previous trial instances is retained, making it mean-centered. Consequently, when prediction errors are minimal, the EV does not substantially change with repeated rewards, rendering the model insensitive to reward frequency.

#### Delta-Prospect-Valence-Learning (Delta-PVL)

The Delta-PVL model (45) extends the basic Delta model by no longer assuming linearity, proportionality, and gain-loss symmetry in the subjective representation of EV, as proposed by the prospect theory (46). Specifically, this model does not assume veridical processing of EV. Instead, it posits that the magnitude of rewards can be transformed by a non-linear shape parameter *γ*, and that individuals may apply different weights to gains versus losses via a gain–loss weighting parameter *λ*. The subjective utility is defined as:

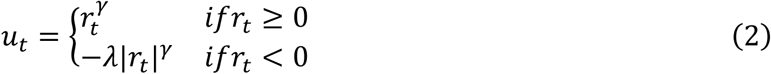

where the shape parameter *γ* (0 < *γ* < 1) determines the curvature of the utility function. When *γ* = 1, all rewards are processed veridically. As *γ* approaches 0, reward magnitudes are increasingly discounted, and at γ = 0, all rewards are treated equivalently (i.e., coded as 1) and the magnitude is completely disregarded. The loss aversion parameter *λ* (0 < *λ* < 5) determines the relative weighting of gains and losses. When *λ* = 1, gains and losses contribute equally; *λ* values below 1 indicate greater sensitivity to gains than losses, whereas *λ* above 1 indicate greater sensitivity to losses than gains.

The computed utility is then entered into the Delta learning rule to update the EV of the chosen option.

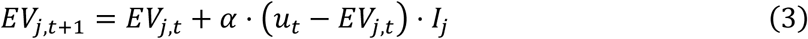

As in equation 1, *I_j_* denotes an indicator for the chosen option and *α* represents the recency learning parameter.

#### Delta-Asymmetric

Because all option EVs in our task were positive and outcomes below zero were rare, we included an additional Delta extension model that applies a relative rather than absolute weighting rule for gains and losses. The Delta-Asymmetric model (47) assigns separate learning rates to positive and negative prediction errors, defining gains and losses based on whether the received reward exceeds or falls short of the expected value for the chosen option. The EVs are then updated using the standard Delta rule:

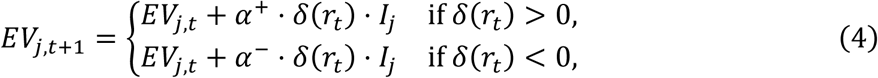

where *δ*(*r*_*t*_) represents the prediction error, *δ*(*r*_*t*_) = *r*_*t*_ − *EV*_*j*,*t*_; *α^+^*and *α^−^* denote the learning rate for positive and negative prediction errors, respectively, *⍺⍺* ∈ (0,1). This model has been shown to effectively account for risk sensitivity in decision-making (47).

#### Decay

The second major class of reinforcement learning rules is the Decay rule (48). In contrast to the Delta rule, which represents a recency-weighted average of reward value or subjective utility, the Decay rule assumes that an option’s EV increases through repeated selection but gradually decays when the option is not chosen. Formally, the Decay rule updates the EV of option *j* as:

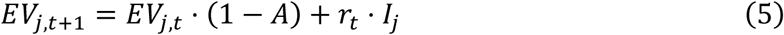

where *A* is the decay parameter akin to *α* in the Delta rule models, *AA* ∈ (0, 1). Higher *A* indicates greater weight assigned to recent outcomes. In this model, the EV for each option gradually decays over time and increases only when a reward for that option is received. As a result, options rewarded more frequently accumulate higher EVs. This mechanism has been shown to capture frequency effects, particularly under conditions where reward frequency may alter participants’ perception of EV (1–3).

#### Decay-PVL

The Decay-PVL model (49) parallels the Delta-PVL model. The conversion from actual rewards to subjective utility follows the same functional form as in Equation 2, but the resulting utility is then integrated using the Decay rule, as follows:

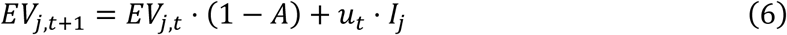

#### Decay-Win

We also fit a relative learning model within the Decay class, adapted from the Prediction-Error Decay model (16). In this Decay-Win model, EV accumulation depends solely on how often an option yields above-average outcomes. Here, rewards are defined relative to whether the obtained outcome exceeds the running average, *AV*, updated as:

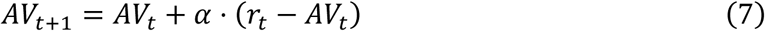

and the EV is calculated as:

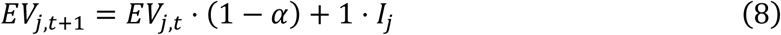

where *I_j_* an indicator term set to 1 if *r_t_* > *AV_t_* and 0 otherwise. In this model, only the valence of the outcome (i.e., whether it is a “win” or not) is used to guide people’s choices. EVs decay over time, and increments occur only when rewards surpass the overall average. The model ignores exact reward magnitudes and instead tracks only the number of above-average outcomes associated with each option, thereby providing a clean dissociation between frequency-based and value-based processing.

#### Hybrid Model

Finally, we selected the best-fitting models from each class to form a hybrid model, consisting of the two relative models: Delta-Asymmetric and Decay-Win (see *Results*). The inclusion of the Decay-Win model as the representative of the Decay rule class completely dissociates value-based processing from frequency-based processing: the Delta component (i.e., Delta-Asymmetric) retains no memory of reward frequency, while the Decay component (i.e., Decay-Win) retains no memory of actual reward values. This hybrid configuration also provided the best model fit among all tested combinations of hybrid models (see *Supplementary Table 2*).

All simple models used a *SoftMax* rule to convert EVs into each model’s predicted probability of selecting each *j* alternative on trial *t*:

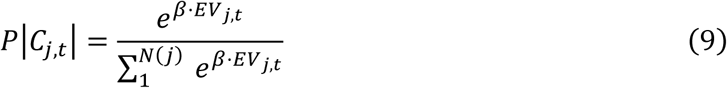

where *β* = 3^*c*^ − 1 (0 ≤ *c* ≤ 5), and *c* is a log inverse temperature parameter that determines how consistently the option with a higher EV is selected (50). When *c* = 0, choices are random; as *c* increases, the option with the highest EV is selected more often. For the hybrid model, a free weighting parameter, *w*, was applied to the choice probabilities generated by each component process and the final predicted probability is calculated as:

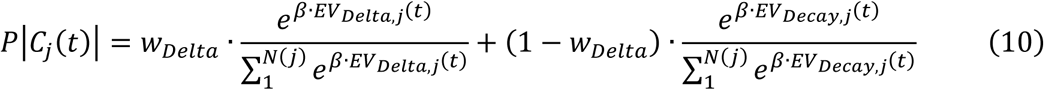

#### Model Fitting and Evaluation

We used the maximum likelihood (ML) approach for model fitting. The negative log likelihood of the parameter set *θ*, given observed data *y* and model 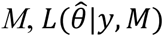, was minimized using the *minimize* function in the *SciPy* library in Python. To avoid local minima, optimization was repeated 100 times with randomly selected starting points for each parameter. All trials except the first trial in each condition were included for model fitting, and the outcome or utility of the first trial in each condition was used to initialize EV values.

Model comparison relied on the Bayesian Information Criterion (BIC) (51). For each participant in each condition, we computed:

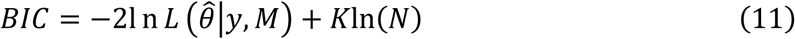

where *N* is the number of observations. In our study, *N* equals 200 trials in each condition. To evaluate model evidence, we calculated BIC weights (52):

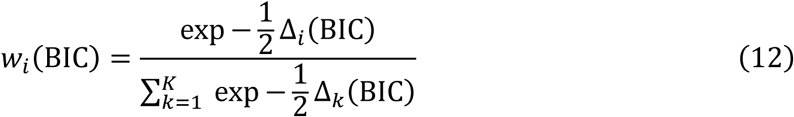

where Δ_*i*_(BIC) = BIC_*i*_ − *m*i*n* (BIC). Lower BIC values indicate better model fit, whereas higher BIC weights reflect stronger relative support for a given model. We also calculated the Bayes Factor (BF_10_) using:

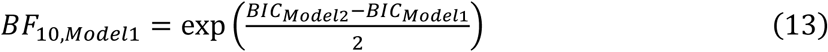

with BF_10_ > 3 generally considered significant, representing a moderate advantage in favor of Model 1 (53).

In addition, we conducted group-level model comparisons using Variational Bayesian Model Selection (VBMS) (54). VBMS treats each model as a random variable and estimates the parameters of a Dirichlet distribution, which are then used to construct a multinomial distribution describing the probability that each model generated the data of a randomly chosen participant. Posterior Dirichlet parameters, *α*, represent the estimated frequency with which each model best explains individual participants’ data. The posterior multinomial parameter, *r_k_*, gives the probability that data from a randomly chosen participants were generated by mode *k*. Finally, the exceedance probability, *φ_k_*, quantifies the likelihood that a particular model *k* is more likely than all competing models to generate group-level data. We used BIC to approximate the log evidence and all three metrics were calculated for model comparisons.

## Results

### Overall Behavioral Results

We first examined participants’ performance during the training phase using a mixed-effects logistic regression model, predicting trial-wise choices from condition, trial type, and block. This analysis revealed a main effect of trial type (*β* = 0.145 ± 0.039, *t* = 3.680, *p* < .001), with participants demonstrating higher accuracy on CD trials than AB trials, and a main effect of block (*β* = 0.039 ± 0.007, *t* = 5.423, *p* < .001), with a greater proportion of optimal choices observed over time. There was no main effect of condition (*β* = 0.056 ± 0.037, *t* = 1.516, *p* = .130), indicating comparable learning effects across conditions.

The only significant interaction was between trial type and condition (*β* = -0.161 ± 0.057, *t* = -2.807, *p* = .005). As shown in *Figure 2a*, the performance gap between AB and CD trials was significantly reduced in the frequency condition. This may reflect either poorer learning due to reduced exposure to CD trials or increased exploratory behavior, as participants may perceive CD trials as scarce opportunities worth exploring. All other interactions—including trial type × block (*β* = 0.008 ± 0.010, *t* = 0.762, *p* = .446), condition × block (*β* = -0.003 ± 0.009, *t* = -0.281, *p* = .779), and the three-way interaction (*β* = 0.009 ± 0.015, *t* = 0.606, *p* = .544), were insignificant.

**Figure 2.**
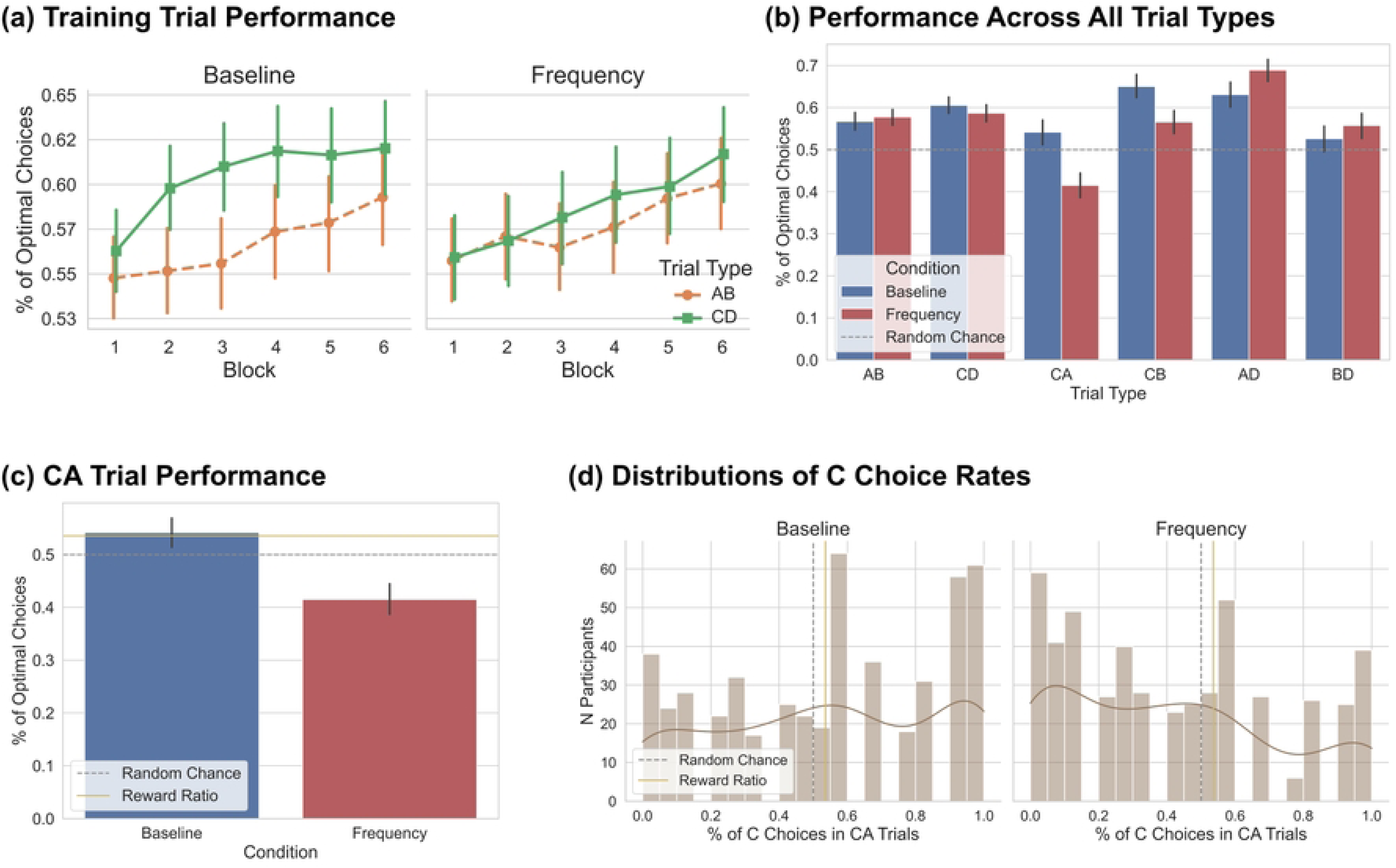
Behavioral Results. *Behavioral Results.* (a) Participants generally showed improved performance across blocks. Overall training accuracy did not differ significantly between conditions, but the typical advantage in CD trials over AB trials observed in the baseline condition disappeared in the frequency condition. (b) Participants selected the optimal option at rates significantly above chance across most trial types, with only two exceptions: BD trials in the baseline condition where the optimal choice rates were not significantly different from chance, and CA trials in the frequency condition, where participants significantly favored the more frequently rewarded, yet suboptimal option A. (c) A closer look at CA trials revealed that in the baseline condition, C choice rates closely matched the objective reward ratio, whereas in the frequency condition they fell significantly below both the reward ratio and chance level. (d) The distribution of C choice rates further confirmed this effect and showed that a larger proportion of participants in the frequency condition chose C less often than either the reward ratio or chance level. Error bars indicate 95% CI interval.

To assess overall performance, we examined the percentage of optimal choices across all six trial types in each condition (*Figure 2b*). For each trial type, we conducted a one-sample *t*-test to compare the observed choice rate against random chance level (0.5) and adjusted for multiple comparisons using the Benjamini–Hochberg procedure (55). After correction, participants selected the optimal option at rates significantly above chance across nearly all trial types across both conditions (see *Supplementary Table 3*), with only two exceptions. In the BD trials under the baseline condition, the proportion of optimal choices (*M* = 0.526) was only numerically above chance, *t*(494) = 1.729, *p_adjusted_* = .084, and in the CA trials under the frequency condition, participants significantly favored the suboptimal option A over the optimal option C (see details below).

#### Critical CA Trials

As predicted, in the critical C-A trials, participants significantly favored the optimal option C in the baseline condition, *t*(494) = 2.824, *p_adjusted_* = .005, but showed a reversed preference for the more frequently rewarded yet less valuable option A in the frequency condition, *t*(494) = -5.903, *p_adjusted_* < .001. This between-condition shift in choice preference was statistically significant based on a paired-sample *t*-test, *t*(494) = 6.007, *p* < .001. To further evaluate this effect, we compared participants’ C choice rates against the underlying reward ratio between options C and A, calculated as 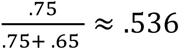. We found that participants’ preference for C (*M* = .541, *SD* = .329) closely matched this expected reward ratio in the baseline condition, *t*(494) = 0.406, *p* = .685, whereas in the frequency condition, C choice rates (*M* = .415, *SD* = .320) fell significantly below this ratio, *t*(494) = -8.388, *p* < .001, indicating a robust reward frequency effect. These results aligned with many prior studies using the same paradigm (1–3), which have consistently demonstrated that, under unequal frequencies of reinforcement, people tend to prefer the more frequently rewarded option even when it yields suboptimal outcomes.

*Figure 2d* shows the distribution of C choice rates across conditions. A larger proportion of participants in the frequency condition chose C less often than both random chance and the reward ratio, further confirming the frequency effect.

#### Within-Subject Results

At the core of the present study lies a key question—whether the seemingly irrational, heuristic-driven frequency effect is more pronounced among better learners and decision-makers. We addressed this question from two angles. First, we examined whether participants with higher training accuracy were more likely to select option A in CA trials under the frequency condition. Second, we tested whether participants who demonstrated better value-based learning performance (i.e., higher C choice rates) in the baseline condition were more likely to choose A in CA trials under the frequency condition.

To address the first question, we ran a series of mixed-effects models predicting the percentage of optimal choices during the testing phase from participants’ combined training accuracy on AB and CD trials, condition, and their interaction. Within the four testing trial types, higher training accuracy was consistently linked with higher optimal choice rates in AD (*β* = 0.507 ± 0.072, *t* = 7.066, *p* < .001) and CB trials (*β* = 0.739 ± 0.072, *t* = 10.287, *p* < .001), and showed no significant relationship in BD trials (*β* = -0.153 ± 0.081, *t* = -1.879, *p* = .061) regardless of condition. However, in CA trials, we found no main effect of training accuracy (*β* = 0.106 ± 0.078, *t* = 1.354, *p* = .176) but a significant interaction effect (*β* = -0.321 ± 0.114, *t* = - 2.828, *p* = .005). In the baseline condition, training accuracy was unrelated—though slightly positively associated—with C choice rates (*β* = 0.106 ± 0.079, *t* = 1.333, *p* = .183), whereas in the frequency condition, the association reversed, with higher training accuracy predicting fewer optimal C choices (*β* = -0.216 ± 0.081, *t* = -2.655, *p* = .008). This reversal was unique to CA trials, as no other trial type showed a change in direction of the training accuracy–optimal choice relationship across conditions (AD: *β* = 0.017 ± 0.104, *t* = 0.162, *p* = .872; CB: *β* = -0.167 ± 0.105, *t* = -1.595, *p* = .111; BD: *β* = 0.080 ± 0.118, *t* = 0.677, *p* = .499; *Figure 3a*). Therefore, we confirmed that while higher training accuracy generally predicted better testing performance, participants with stronger training performance were more likely to choose option A in the frequency condition.

**Figure 3.**
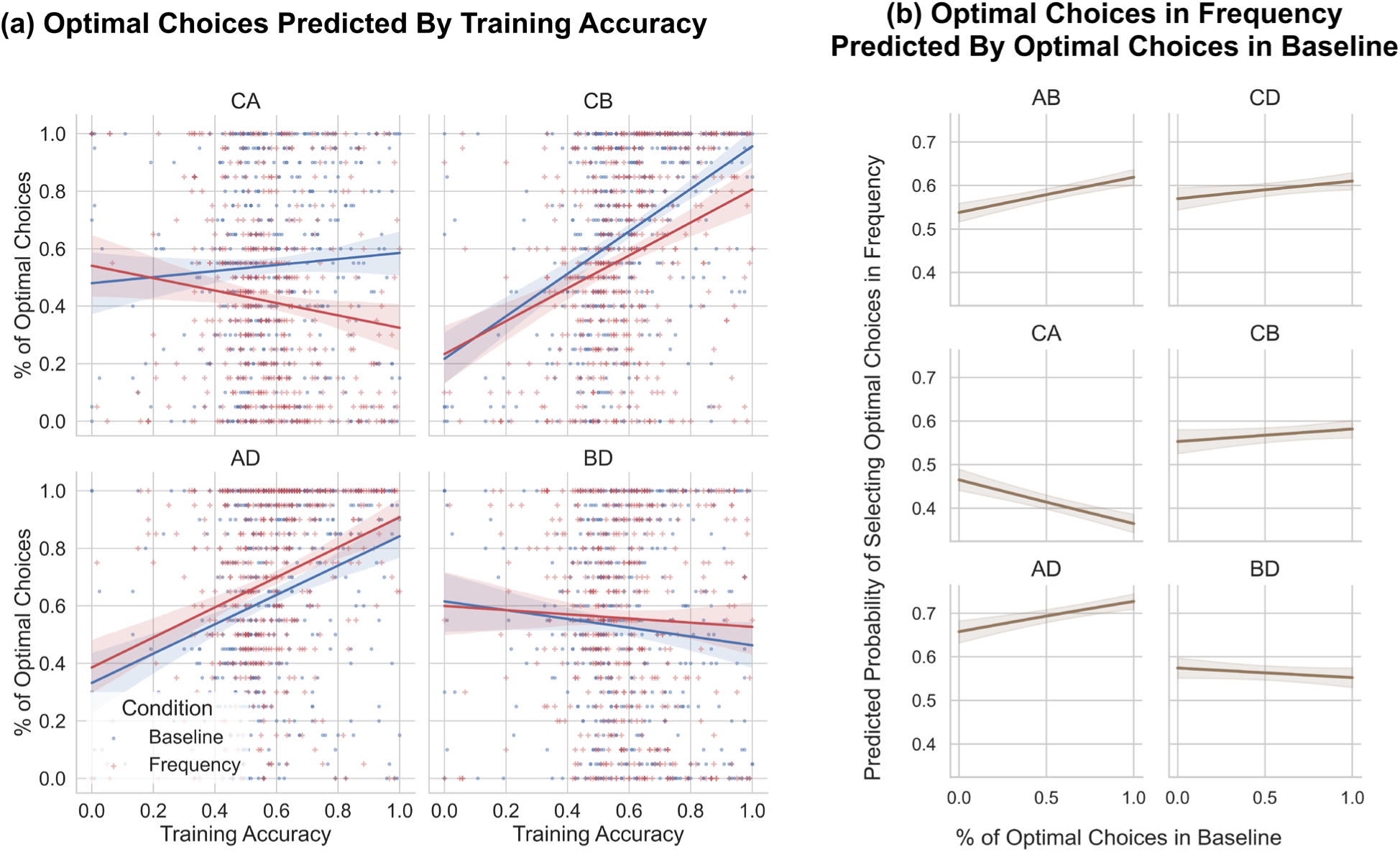
Within-Subject Results. *Within-Subject Results.* (a) For CB and AD testing trials, higher training accuracy was consistently associated with higher optimal choice rates in the testing phase. This association was insignificant for BD trials in both conditions. In contrast, CA trials revealed a significant interaction, where training accuracy was positively (insignificant) related to C choice rates in the baseline condition but negatively related in the frequency condition. This suggests that although better training performance generally promoted optimal test-phase choices, it led participants to show stronger frequency effects when reward frequency was manipulated. (b) Across AB, CD, CB, and AD trials, optimal choice rates were positively correlated between the two conditions, indicating consistent decision-making patterns. For CA trials, however, this association was nonsignificant and numerically negative. Error bars indicate 95% CI interval.

Next, we ran a series of mixed-effects logistic regression models predicting the probability of selecting the optimal option in the frequency condition as a function of the corresponding proportion of optimal choices in the baseline condition, both overall and separately for trial type. Overall, participants’ baseline performance significantly predicted their performance in the frequency condition after controlling for trial types (*β* = 0.061 ± 0.029, *t* = 2.111, *p* = .035), suggesting behavioral consistency across conditions. For specific trial types, this positive association held for AB (*β* = 0.572 ± 0.258, *t* = 2.213, *p* = .027), CD (*β* = 0.606 ± 0.245, *t* = 2.477, *p* = .013), AD (*β* = 0.774 ± 0.342, *t* = 2.263, *p* = .024), CB (*β* = 0.777 ± 0.329, *t* = 2.357, *p* = .018) trials, but not for CA (*β* = -0.412 ± 0.313, *t* = -1.315, *p* = .188) or BD trials (*β* = -0.155 ± 0.344, *t* = -0.449, *p* = .653)—both of which involved closely valued options with different reward frequencies during training. Critically, the slope for CA trials was significantly different—and notably negative—compared to AB (*β* = -0.747 ± 0.091, *t* = -8.175, *p* < .001), CD (*β* = -0.582 ± 0.098, *t* = -5.945, *p* < .001), CB (*β* = -0.532 ± 0.096, *t* = -5.542, *p* < .001), BD (*β* = -0.325 ± 0.096, *t* = -3.372, *p* < .001), and AD (*β* = -0.742 ± 0.103, *t* = -7.238, *p* < .001) trials (*Figure 3b*). These findings suggest that while participants’ selection of optimal choices was overall consistent across conditions, their choice of C in CA trials was uncorrelated, if not negatively correlated, between baseline and frequency conditions. In other words, participants who strongly preferred the optimal option C under baseline conditions did not maintain that preference when option A was more frequently rewarded, suggesting a strategic shift away from value-based decision-making under frequency manipulation.

#### Model Fitting Results

Finally, we applied computational modeling to quantify the relative contributions of reward frequency-based and value-based strategies in each condition. As shown in *Table 1*, the Decay-Win and Hybrid models consistently provided the best fits across participants. The Hybrid model yielded the best average fit as indexed by BIC, while the Decay-Win model accounted for the largest number of individual best fits. The overall advantage of Decay-class models over Delta-class models suggests that participants relied on reward frequency during learning in both conditions. In particular, the superior performance of the Decay-Win model indicates that participants may have relied strongly on binary reward/non-reward tallies.

**Table 1.** Model Fitting Results.

| | $c$ | $\alpha/A/\alpha\text{-pos}$ | $\alpha\text{-neg}$ | $\gamma$ | $\lambda$ | $w$ | BIC<sub>avg</sub> | BIC<sub>weight</sub> | BF<sub>10</sub> | $N_{\text{best fit}}$ | VB $\alpha$ | VB $r_k$ | VB $\phi_k$ |
| --- | --- | --- | --- | --- | --- | --- | --- | --- | --- | --- | --- | --- | --- |
| **Baseline** |  |  |  |  |  |  |  |  |  |  |  |  |  |
| Delta | 1.708 | 0.287 |  |  |  |  | 252.002 | <.001 | 5316.431 | 94 | 54.748 | 0.109 | <.001 |
| Delta PVL | 1.769 | 0.302 |  | 0.548 | 1.862 |  | 256.015 | <.001 | 39527.978 | 12 | 4.597 | 0.009 | <.001 |
| Delta Asymmetric | 1.686 | 0.352 | 0.261 |  |  |  | 242.478 | 0.018 | 45.449 | 43 | 35.244 | 0.070 | <.001 |
| Decay | 0.356 | 0.211 |  |  |  |  | 240.180 | 0.056 | 14.404 | 50 | 44.905 | 0.089 | <.001 |
| Decay PVL | 0.345 | 0.193 |  | 0.357 | 2.118 |  | 242.420 | 0.018 | 44.154 | 30 | 17.254 | 0.034 | <.001 |
| Decay Win | 0.423 | 0.201 |  |  |  |  | 239.151 | 0.094 | 8.611 | 136 | **183.276** | **0.365** | **.874** |
| Hybrid | 1.698 | 0.237 | 0.365 |  |  | 0.528 | **234.845** | **0.813** | - | 130 | 161.976 | 0.323 | .126 |
| **Frequency** |  |  |  |  |  |  |  |  |  |  |  |  |  |
| Delta | 1.766 | 0.299 |  |  |  |  | 253.251 | <.001 | 5220.613 | 98 | 69.907 | 0.139 | <.001 |
| Delta PVL | 1.808 | 0.332 |  | 0.562 | 2.001 |  | 256.807 | <.001 | 30897.749 | 20 | 13.673 | 0.027 | <.001 |
| Delta Asymmetric | 1.796 | 0.382 | 0.240 |  |  |  | 245.337 | 0.008 | 99.864 | 34 | 18.770 | 0.037 | <.001 |
| Decay | 0.379 | 0.225 |  |  |  |  | 241.835 | 0.048 | 17.336 | 75 | 56.601 | 0.113 | <.001 |
| Decay PVL | 0.374 | 0.221 |  | 0.340 | 2.084 |  | 244.168 | 0.015 | 55.648 | 21 | 16.873 | 0.034 | <.001 |
| Decay Win | 0.416 | 0.195 |  |  |  |  | 240.334 | 0.101 | 8.185 | 129 | **189.481** | **0.377** | **.998** |
| Hybrid | 1.664 | 0.274 | 0.381 |  |  | 0.517 | **236.130** | **0.828** | - | 118 | 136.695 | 0.272 | .002 |

#### Parameter Analysis

To further examine within-subject changes in strategy use and identify individuals who exhibit stronger frequency effects, we analyzed whether participants’ baseline behavior could predict reliance on frequency-based strategies in the frequency condition. We first focused on simpler, single-process models. A mixed-effects model was conducted to predict model fit (i.e., BIC) of the three Decay rule models in the frequency condition with participants’ C choice rates in the baseline condition. We controlled for model type to reduce model-specific noise. Results revealed that higher C choice rates in the baseline condition significantly predicted better fit of Decay rule models in the frequency condition (*β* = -17.699 ± 6.939, *t* = -2.551, *p* = .011), suggesting that individuals who learned value-based contingencies well in baseline condition were more likely to be better captured by frequency-sensitive Decay rule models when reward frequency was manipulated (*Figure 4a*). Crucially, this relationship did not hold for frequency-insensitive Delta rule models (*β* = -9.632 ± 6.353, *t* = -1.516, *p* = .130), whose fit was unrelated to participants’ baseline C choice rates.

**Figure 4.**
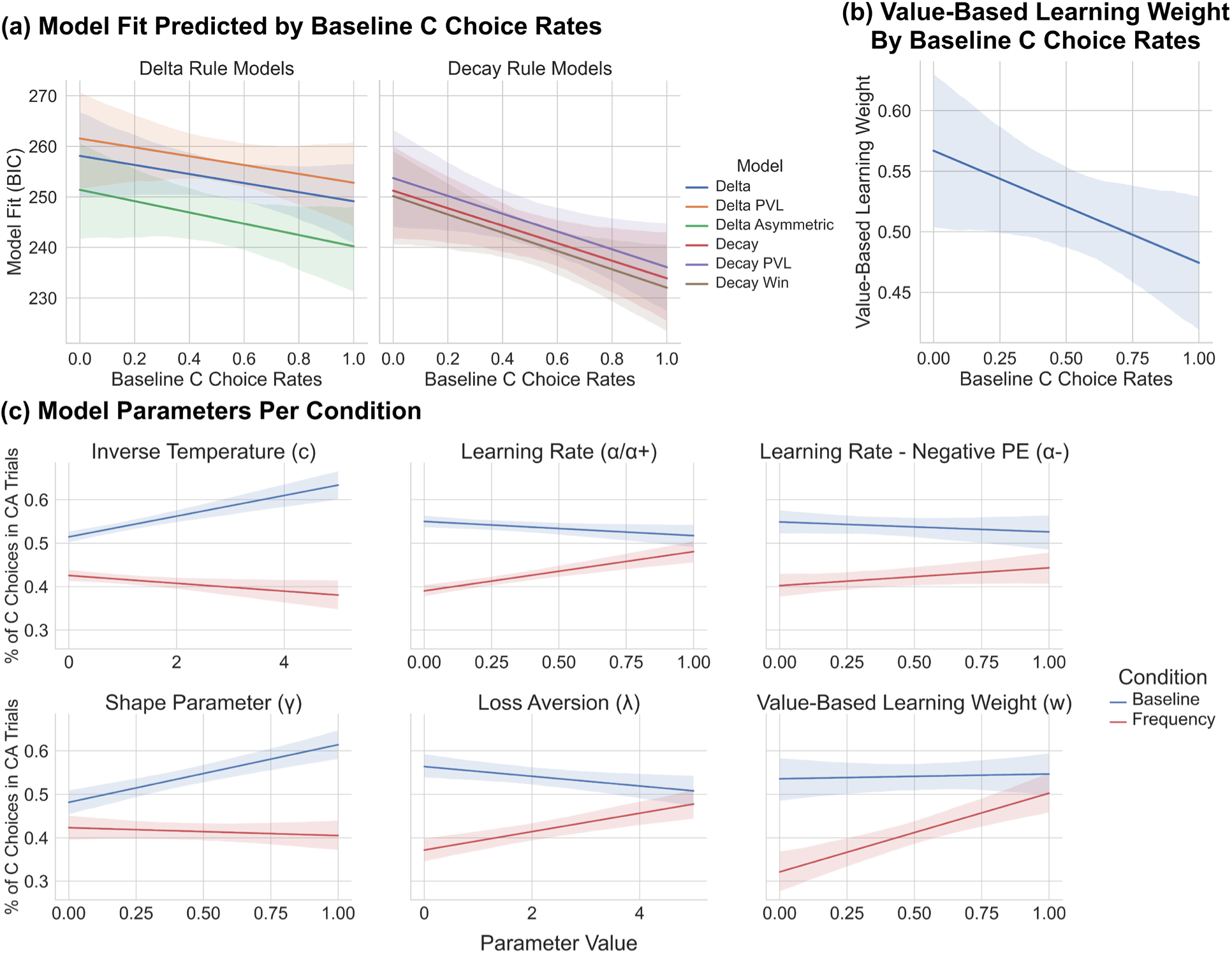
Model Fitting Results. *Model Fitting Results.* (a) Higher C choice rates in the baseline condition significantly predicted better fit for Decay-class models in the frequency condition, but not for Delta-class models. (b) Examination of the weight parameter in the Hybrid model showed that higher baseline C choice rates predicted reduced reliance on value-based Delta processing in the frequency condition. This suggests that individuals with higher baseline accuracy in CA trials may absorb less information from value-based strategies and rely more on frequency-based processing when reward frequency was manipulated. (c) All model parameters showed significant interaction effects across the two conditions, such that parameters positively associated with C choice rates in the baseline condition predicted the opposite or null effect in the frequency condition, and vice versa. Given the strong parameter correlations across conditions, this pattern suggests that traits leading individuals to favor C in the baseline condition may also predispose them to shift toward A in the frequency condition. Error bars indicate 95% CI interval.

Next, we examined the hybrid model to assess whether participants who selected option C more frequently in the baseline condition were more likely to shift toward a frequency-based strategy under frequency manipulation. Specifically, we ran a general linear model predicting the model-inferred Delta weights (i.e., the relative reliance on value-based learning) in the frequency condition based on participants’ C choice rates in the baseline condition, controlling for training performance. The results revealed a significant negative correlation: participants who chose C more often in the baseline condition showed lower Delta weights in the frequency condition (*β* = -0.103 ± 0.052, *t* = -1.977, *p* = .049), indicating greater reliance on the frequency-based process (*Figure 4b*). Importantly, this pattern was consistent across alternative, less well-fitting hybrid configurations where different Delta and Decay variants were combined (*Supplementary Figure 2*). Together, these findings suggest that individuals who previously engaged in more rational, value-based learning were particularly susceptible to switching to a heuristic, frequency-driven strategy when reward frequencies were unequal and demonstrating stronger frequency effects.

Finally, we looked into the best-fitting model parameters. Parameter estimates were strongly correlated across conditions (*β* = 0.158 ± 0.010, *t* = 15.935, *p* < .001) after controlling for parameter type and model type, indicating high consistency in general decision-making characteristics across conditions. However, mixed-effects models predicting the percentage of C choices in CA trials based on condition and specific parameters revealed significant interactions across all parameters (*c*: *β* = -0.031 ± 0.004, *t* = -7.451, *p* < .001; *α*: *β* = 0.099 ± 0.018, *t* = 5.560, *p* < .001; *α-neg*: *β* = 0.075 ± 0.035, *t* = 2.128, *p* = .034; *γ*: *β* = -0.136 ± 0.031, *t* = -4.432, *p* < .001; *λ*: *β* = 0.030 ± 0.006, *t* = 5.230, *p* < .001; *w*: *β* = 0.170 ± 0.054, *t* = 3.164, *p* = .002). Specifically, parameters associated with higher C choice rates in the baseline condition were associated with lower C choice rates in the frequency condition, and vice versa (*Figure 4c*). Given the overall stability of individuals’ best-fitting parameters across conditions, these results indicate that individuals whose traits inclined them to favor C in the baseline condition are more likely to shift towards A in the frequency condition.

## Discussion

In the science of decision-making, it has long been argued that the use of heuristic, fast-and-frugal strategies may be adaptive (4,11,33,34). Yet no study has systematically examined whether such heuristics are more or less likely to emerge in individuals who demonstrate stronger learning performance. In the present study, we addressed this question using a within-subject design in which each participant completed the task twice—once under baseline conditions and once with a reward-frequency manipulation in an environment known to elicit frequency effects (2,3).

Our manipulation successfully altered participants’ behavior. Participants strongly preferred the more valuable option C in the baseline condition but shifted to favor the less valuable option A when A was presented twice as often during training in the frequency condition. Within-subject analyses further revealed two key patterns. First, participants with better training performance generally achieved higher accuracy in the testing phase, except in CA trials under the frequency condition, where higher training accuracy predicted a stronger tendency to favor the frequently rewarded but less valuable option A. Second, accuracy across trial types was generally consistent between conditions, whereas in the critical CA trials, C choice rates in the baseline condition did not predict, and even negatively trended with, C choice rates in the frequency condition.

Computational modeling validated these behavioral findings. Participants with higher baseline C choice rates were found to be better fit by frequency-sensitive Decay class models in the frequency condition but not frequency-insensitive Delta class models. In the Hybrid model, which combines a purely value-based and a purely frequency-based process with a weighting parameter, these good learners showed greater reliance on the frequency-based process when the less valuable option was rewarded more often. Together, these results suggest that individuals who demonstrate stronger baseline learning are also those more likely to shift toward frequency-based processing when strong frequency cues are present, thereby exhibiting stronger frequency effects.

Since the IGT gained popularity, it has often been assumed that focusing on reward frequency while neglecting underlying reward values reflects impaired value learning associated with neuropsychological deficits (14,15,56,57). However, more recent studies consistently demonstrate that even healthy participants prefer options yielding more frequent rewards, despite their lower average value (2,3,21–24,30). This suggests that reward frequency is not merely a marker of deficit, but potentially a fundamental component of human value learning and decision-making. Evidence from both the IGT and its inverted version further underscores this point by demonstrating that frequent small wins outweigh infrequent large losses, leading participants to view the frequently rewarded option as superior (21,23,58). Conversely, frequent small losses overshadow infrequent large wins, prompting participants to avoid these options even when the long-term value is positive (59). In the present study, we extend this literature by showing that reward frequency plays an indispensable role in shaping subjectively perceived values. Crucially, individuals with stronger baseline learning and decision-making performance exhibited more pronounced, seemingly irrational, frequency effects. This pattern supports the view that frequency effects are not simply a sign of flawed learning in complex environments. Instead, they may reflect an adaptive strategy shift, where good learners are able to actively perceive, process, and integrate frequency cues into their decision-making as part of a flexible response to environmental demands.

That said, our findings do not preclude the possibility that a rigid, excessive focus on reward frequency can be maladaptive. Prior research shows that frequency effects at the group level typically emerge only under conditions of highly elevated environmental uncertainty (3). Attending exclusively to immediate rewards, without regard to long-term payoffs, in environments that do not necessitate such a strategy switch may signal deficient cognitive functioning, such as low self-control (60). Thus, the adaptiveness of seemingly irrational heuristics, such as frequency effects, depends on whether the environment calls for a flexible shift away from purely value-based decision-making, but may become harmful when value-based strategies remain tractable. Overall, our findings emphasize that frequency effects are neither a mere cognitive flaw nor a one-size-fits-all strategy. Rather, it may serve as a context-dependent adaptive tool that enables decision-makers to flexibly balance efficiency and accuracy when navigating uncertain environments.

One limitation of the present study is that adaptive switching from value-based decision-making to seemingly irrational heuristics was examined only in the context of frequency effects in reinforcement learning. Other heuristics that are not necessarily tied to reward frequency— such as take-the-best, one-clever-clue, and fast-and-frugal trees (4)—require further investigation to determine whether they, too, emerge more strongly in better learners. In addition, it remains unclear whether such strategy switching persists in environments characterized by low to moderate uncertainty. We speculate that individuals with stronger learning performance may be more resilient to frequency-based biases when the environment is relatively simple to grasp, but this possibility awaits empirical testing.

In conclusion, the present study demonstrates that frequency effects, as long considered markers of impaired value learning, may instead reflect an adaptive shift in strategy, particularly among individuals with stronger learning performance. By combining behavioral analyses with computational modeling, we showed that better learners not only displayed stronger frequency effects under reward-frequency manipulation but were also best captured by frequency-sensitive models. These findings suggest that what appears to be irrational bias may, under conditions of uncertainty, represent a flexible adaptation when value-based strategies are costly or unreliable. Ultimately, clarifying when and why individuals adopt heuristic strategies will advance our understanding of the balance between rational value learning and adaptive shortcuts in human decision-making, offering deeper insight into the nature of human rationality in ecological, real-world contexts.

## Acknowledgement

None.

